# Robo1 and 2 repellent receptors cooperate to guide facial neuron cell migration and axon projections in the embryonic mouse hindbrain

**DOI:** 10.1101/294702

**Authors:** Hannah N. Gruner, Minkyung Kim, Grant S. Mastick

## Abstract

**Background:** The facial nerve is necessary for our ability to eat, speak, and make facial expressions. Both the axons and cell bodies of the facial nerve undergo a complex embryonic migration pattern involving migration of the cell bodies caudally and tangentially through rhombomeres, and simultaneously the axons projecting to exit the hindbrain to form the facial nerve.

**Results:** Our goal in this study was to test the functions of the chemorepulsive receptors Robo1 and Robo2 in facial neuron and axon migration by analyzing genetically marked motor neurons in double mutant mouse embryos through the migration time course, E10.0-E13.5. In Robo1/2 double mutants, axon and cell body migration errors were more severe than in single mutants. Most axons did not make it to their motor exit point, and instead projected into and longitudinally within the floor plate. Surprisingly, some facial neurons had bifurcated axons that either exited or projected into the floor plate. At the same time, a subset of mutant facial cell bodies failed to migrate caudally, and instead shifted into the floor plate.

**Conclusions:** Robo1 and Robo2 have redundant functions to guide multiple aspects of the complex cell migration of the facial nucleus, as well as regulating axon trajectories and suppressing formation of ectopic axons.

**Bullet points:**

1. Robo1 and 2 repellent receptors were tested for functions in facial neuron cell body and axon migration in the developing E10.0-E13.5 hindbrain of double mutant mouse embryos.
2. In Robo1;Robo2 double mutant embryos, facial motor neuron cell bodies and axons abnormally grew into the hindbrain floor plate, with stronger effects than in single Robo mutants.
3. Many facial nerve axons abnormally bifurcated, failed to project to their proper exit point, and instead projected longitudinally within the floor plate.
4. Facial neuron cell body tangential migration was disrupted.
5. Robo receptor-mediated repulsion is required for proper migration of both facial axon projections and cell body migrations.

Funding was provided to G.S.M. by NIH R01 NS054740, R21 NS077169, and R01 EY025205. Core facilities at the University of Nevada Reno campus are supported by NIH COBREs GM103650 and GM103554, and Nevada INBRE GM103440.

## Introduction

In the developing nervous system, neural progenitors must accurately navigate through the complex environment of the embryo to arrive at their final target. Neurons differentiate in specific locations and send out axon projections that are directed by a combination of guidance cues (Tessier-Lavigne et al., 1988). In the hindbrain, the rhombomere segments compartmentalize the differentiation of many of the cranial nerve populations, including the facial nerve (Lumsden and Keynes, 1989). From their birthplace in rhombomere 4 (r4), the facial branchiomotor nucleus (FBMN) undergoes a complex series of neuron cell body migrations while simultaneously sending out axons to exit to form the facial nerve (Auclair et al., 1996; Fritzsch, 1998; Fritzsch and Nichols, 1993). However, key aspects of the guidance mechanisms for how the cell bodies and axons migrate in the CNS remain unclear.

The intricate migratory pattern of the facial nucleus and nerve suggests that integration of multiple guidance pathways is necessary to guide this population. In mouse embryos, cell bodies of the branchiomotor (BM) aspect of the facial nucleus first differentiate in r4 via Hoxb1, Islet-1, Phox2b, and hedgehog activity (Chandrasekhar et al., 1998; Goddard et al., 1996; Kim et al., 2016; Pattyn et al., 2000; Studer et al., 1996; Varela-Echavarría et al., 1996; Zhuang et al., 2013). After appearing in ventral r4 on embryonic day 9.5 (E9.5), facial axons begin to arrive by E10.0 at their motor exit point in dorsal r4 (Niederländer and Lumsden, 1996). These initial projections by motor neurons are thought to be guided by an unknown attractive cue secreted by the exit point, as well as repulsive cues restricting where axons can migrate (Bravo-Ambrosio and Kaprielian, 2011; Bravo-Ambrosio et al., 2012; Guthrie and Lumsden, 1992; Xiao et al., 2003). During E10.5-E12.5, the chemorepulsive receptor Neuropilin-1 (Nrp1) and its ligands Sema-3A and VEGF164 are required for tangential migration and distal fasciculation of the facial cell bodies and axons, respectively (Kawakami et al., 1996; Kitsukawa et al., 1997; Schwarz et al., 2004; Varela-Echavarría et al., 1997).

While the facial axons exit to begin their navigation to targets in the periphery, their cell bodies are also undergoing complex migrations in the hindbrain. Beginning on E10.5, cell bodies shift from their ventral position in r4 to a more caudal location in r6. This translocation occurs via tangential migration, in which cell bodies translocate behind small leading axon-like protrusions that direct the cell body migration. Members of the Polar Cell Polarity (PCP) pathway (Prickle1, Prickle1b, Vangl2, Protein tyrosine kinase 7, Celsr1-1a-2-3, Frizzled3a, Frizzled7, Tbx20, Wnt5a, Scribble) are required to mediate this first step of caudal migration in mouse and in zebrafish (Carreira-Barbosa et al., 2003; Glasco et al., 2012; Mapp et al., 2010; Qu et al., 2010; Song et al., 2006; Vivancos et al., 2009; Yang et al., 2014). Mutating these genes prevents or limits facial cell bodies from successfully translocating to r6, implying these are necessary to promote caudal migration. Interestingly, the PCP proteins appear to act independently of Dvl1/2, suggesting a unique signaling cascade that utilizes different signaling molecules than most known Wnt pathways (Glasco et al., 2012). This indicates that other effectors, perhaps activated by Robo-Slit signaling, are utilized in the facial nucleus to activate Rho GTPAse mediated cytoskeletal rearrangements.

After arriving in r6, the facial neurons further migrate within r6, with ventral to dorsal migrations, and then ventricular to pial. These shifts are not as well defined mechanistically, but initiate in r6 on E12.5 and finish at E14.5. One factor identified for the dorsolateral migration is the transcription factor Ebf1, as knockouts exhibit premature dorsolateral migration of the FBMN into r5 (Garel et al., 2000). The Dachsous-Fat PCP pathway is also necessary for dorsolateral migration, as Fat4/Dchs1 mutants fail to initiate this migration, and instead remain next to the floor plate (Zakaria et al., 2014). As the dorsal translocation is occurring, the cells also migrate radially from the ventricular surface to the pial surface. In Reeler mutant mice, the facial nuclei become disorganized and do not reach the pial surface, suggesting this migration depends on radial glial substrates (Goffinet, 1984; Rossel et al., 2005). Similarly, Cdk5, Dab1, and p35 are necessary for facial cell bodies to migrate to the pial surface (Ohshima et al., 2002). Although much is known about the migration of these neurons, our current knowledge paints an incomplete picture of which external signaling pathways are involved. In addition, the extent to which cell body and axonal migration are separable or coupled events is unclear.

Slit-Robo repellents are highly conserved guidance molecules necessary to guide many populations of cells during development (Brose et al., 1999; Kidd et al., 1998; Zallen et al., 1998). This signaling pathway guides a multitude of neuronal populations (Dickson and Gilestro, 2006; Mastick et al., 2010; Ypsilanti et al., 2010), and in a relatively few number of cases to guide cranial nerves (Cho et al., 2007; Fouquet et al., 2007; Li et al., 1999; Prince et al., 2009).

A previous study identified Slit-Robo functions to guide facial axons (Hammond et al., 2005). First, they established that the FBMN express Robo1/2 early (E12.0 rat, equivalent to E10.0 mouse), while their ligands Slit1 and 2 are expressed by the flanking ventral floor plate and dorsal rhombic lip (Hammond et al., 2005). This study then examined mouse embryos by retrograde labeling from the facial nerve at one early stage, E11.5, and found that single Robo mutants, particularly Robo2, had most axons projecting to exit normally while a few FBMN axons entered the floor plate. Slit-Robo signaling was shown to organize and guide facial axons toward the dorsal motor exit point at r4, while also repelling axons out of the Slit-producing floor plate, results which were supported in chick embryos using dominant negative constructs. Robo2 was found to be of particular importance for restricting facial axons to r4, preventing a subset of axons from coursing rostral or caudal relative to r4. How Robo-Slit signaling influences the facial nerve and nucleus at earlier or later stages, and particularly the potential redundant functions of Robo1 and 2 in FMBN development, remained unresolved.

To extend the prior study, we set out to more closely examine the role of Robo-Slit signaling during the development of the facial nerve and nucleus by 1) using multiple time points during development, 2) analyzing FBMN defects in Robo1/2 double mutant mice, and 3) using genetic markers for FBMN neurons in addition to retrograde tracing. We found that Slit-Robo signaling guides and organizes facial axons throughout several stages of neuron migration and axon projection. Additionally, we found enhanced defects in the positioning of FBMN cell bodies in Robo1/2 mutant embryos. Our data suggests a complex guidance mechanism by which a population of neuronal cell bodies and their axons simultaneously use Robo signaling in several steps of migration in the hindbrain.

## Results

### Isl1-EGFP transgenic mice to visualize Robo-dependent FBMN axon and cell body migration

Our overall goal was to compare the migration of the facial nucleus throughout development between control and Robo1/2 double mutant embryos. To map the development of the facial branchiomotor neurons, we used transgenic Isl^MN^:GFP-F embryos during various stages of development. This transgenic line expresses EGFP in motor neurons under the control of the Isl1 enhancer, crest1 (Lewcock et al., 2007), and proved to be effective to study the development of the FBMN. We first sought to confirm and extend the results of Hammond et al., which relied on retrograde tracing from the nerve exit point, at a single early stage, E11.5, with defects in axon projections seen *Robo1*^+/+^; *Robo2*^−/−^ embryos (Hammond et al., 2005). Their retrograde tracing strategy would only label neurons which successfully projected axons to the exit point, whereas the Isl^MN^;GFP transgenic marker would label all motor neurons and their axons, even those that failed to project an axon to the exit.

In control Isl^MN^;GFP embryos, we observed the expected morphology of the FBMN at E11.5, consisting of many fascicles extending towards the dorsal motor exit point at r4 (Figure 1A, D) and corresponding cell bodies transiting from r4 to r6 (Figure 1A-C). The neuronal cell bodies of the FBMN were located in streams, extending from r4 to r6 on either side of the floor plate, and as expected in control embryos, with no neuronal cell bodies located in the floor plate. As previously reported, there were few longitudinal axons within the floor plate of control embryos (Figure 1B, C) (Hammond et al., 2005).

**Figure 1.**
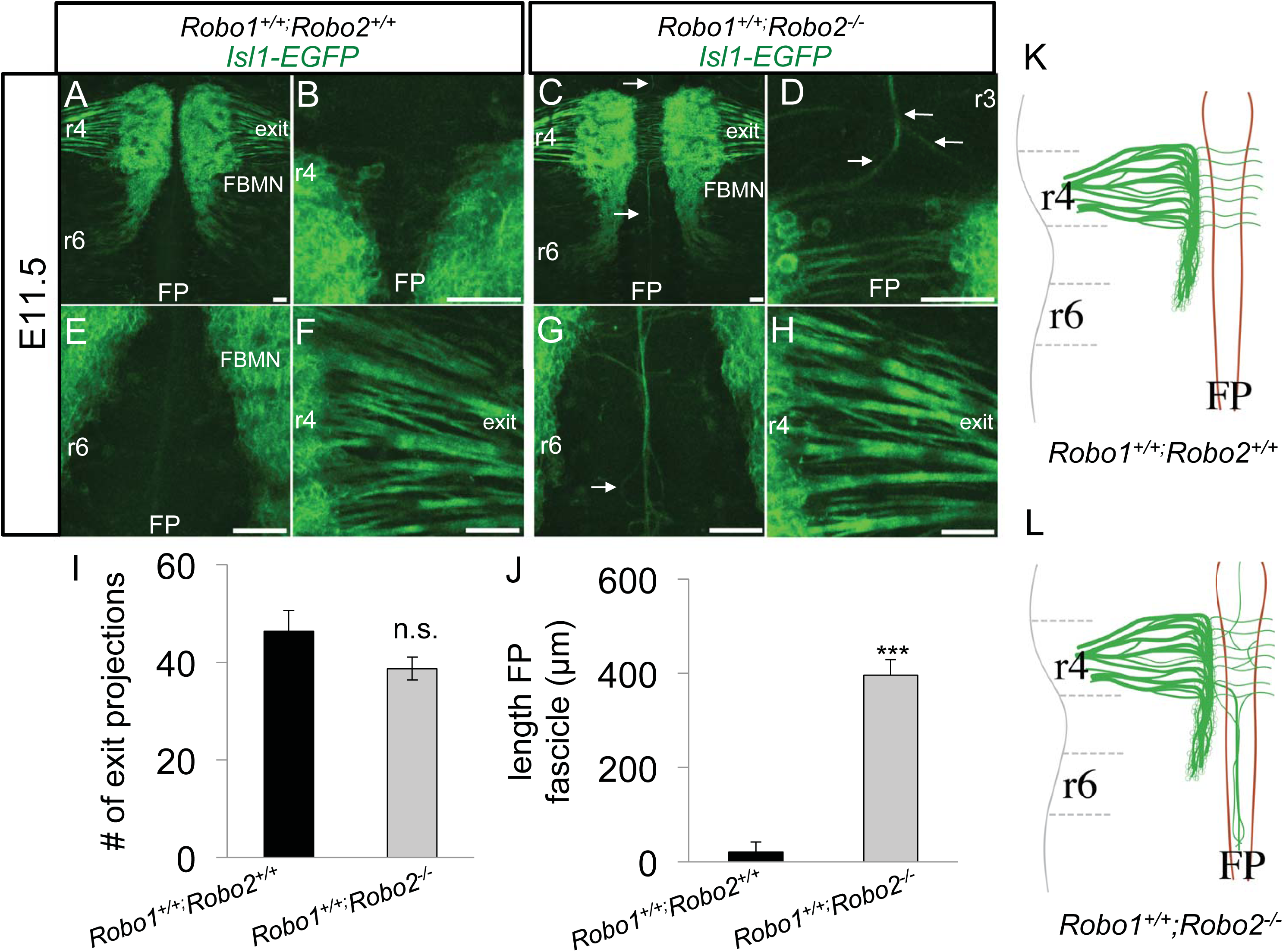
A subset of facial axons enter the hindbrain floor plate in Robo2 single mutants. Transgenic mice, Isl^MN^:GFP, expressing EGFP under the control of somatic motor neuron specific enhancer, Isl1, were utilized to visualize the facial nucleus and nerve. E11.5 open-book Isl1-EGFP hindbrains comparing control **(A-D)** and *Robo1*^+/+^;*Robo2*^−/−^ **(E-H)** embryos. **(A, D)** In control embryos, axons from the FBMN successfully migrated towards the dorsal motor exit point (denoted as exit) in r4; a higher magnification view is shown in panel D. **(A-C)** Control embryos had no apparent longitudinal axons within the r3 (B) or r6 (C) floor plate. **(E, H)** Similar to control, the majority of *Robo1*^+/+^;*Robo2*^−/−^ FBMN axons migrated successfully to their r4 exit point embryos. **(E-G)** FBMN axons projected longitudinally into the floor plate (arrows), migrating to r3 (F) and r6 (G). **(I)** Quantification of the number of fascicles branching directly off the FBMN nucleus. **(J)** Average length of fascicles migrating into the caudal FP in control and *Robo1*^+/+^;*Robo2*^−/−^ embryos. **(K, L)** Schematics of E11.5 FBMN nucleus and axon migration patterns. **(K)** In control animals, axons projected to the exit point in rhombomere 4. **(L)** In Robo2 mutants, some axons collapsed into the floor plate. Scale 50 μm. n=3, E11.5 *Robo1*^+/+^;*Robo2*^+/+^. n=3, E11.5 *Robo1*^+/+^;*Robo2*^−/−.^

In *Robo1*^+/+^; *Robo2*^−/−^ E11.5 littermate embryos carrying the Isl^MN^;GFP, marker, the organization of the FBMN was somewhat altered, in agreement with Hammond, et al.. We saw longitudinal axons projecting from the FBMN migrating into the floor plate (Figure 1E-G). Axons of the FBMN migrated into the floor plate both rostral (Figure 1F) and caudal (Figure 1G) relative to r4. Longitudinal axons in the floor plate emanating from the FBMN largely formed one large fascicle, however some axons appeared to defasciculate and then re-associate with the main bundle (Figure 1G). The majority of *Robo1*^+/+^;*Robo2*^−/−^ axons appeared to successfully fasciculate and correctly exit at r4. Comparing the Robo2 mutants to controls, there was no statistically significant reduction in the number of fascicles emanating from the FBMN going towards the correct r4 exit point (Figure 1H, I). Interestingly, though there was no statistical reduction in the number of axons projecting towards the exit point, there was a dramatic increase in the length of caudally projecting axons found in the floor plate (Figure 1J). Together these data demonstrate the effectiveness of using Isl^MN^;GFP to study the development of the FBMN. Our observations also confirm the role of Robo2 preventing the entry of some FBMN axons into the floor plate (Hammond et al., 2005).

Next we sought to examine if Robo1 and Robo2 play redundant roles in guiding the FBMN. Though Robo2 mutants appear to show some disorganization of FBMN axons, while Robo1 mutants have normal FBMN patterns (Hammond et al., 2005), it is possible that Robo2 may mask functions of Robo1 in guiding the FBMN. To start teasing apart the possible redundant functions of Robo1 and 2, *Robo1*^*+/−*^; *Robo2*^*−/−*^ E11.5 Isl^MN^;GFP embryos were compared to control littermates.

Again, control embryos showed the conventional morphology (Figure 2A-D). There were no longitudinal axons observed in the r3 floor plate (Figure 2B). The cell bodies migrated through r5 towards r6 (Figure 2C), and axonal projections in r4 successfully reached the motor exit point (Figure 2D). *Robo1*^+/−^; *Robo2*^−/−^ embryos exhibited overall normal organization of the FBMN nucleus, suggesting that one allele of Robo1 is largely sufficient for guidance of the cell body migration (Figure 2E). More pronounced than *Robo1*^+/+^; *Robo2*^−/−^ animals, *Robo1*^+/−^; *Robo2*^−/−^ mutants had increased axons migrating into the floor plate (Figure 2F, G). In agreement with this observation, there were somewhat fewer FBMN fascicles projecting towards the r4 exit point in *Robo1*^+/−^; *Robo2*^−/−^ embryos (27% of controls) (Figure 2H, I), suggesting that some FBMN axons projected into the floor plate instead of their correct route. Together these data indicate Robo1 and Robo2 indeed function redundantly in guiding the FBMN axons away from the floor plate. Despite the moderately severe axon guidance errors observed, there was no difference in the tangential caudal migration of the cell bodies **(J)**. Together, the increased errors in *Robo1*^+/−^; *Robo2*^−/−^ mutants compared to *Robo1*^+/+^; *Robo2*^−/−^ suggest that Robo1 and Robo2 have redundant functions in guiding the FBMN axons.

**Figure 2.**
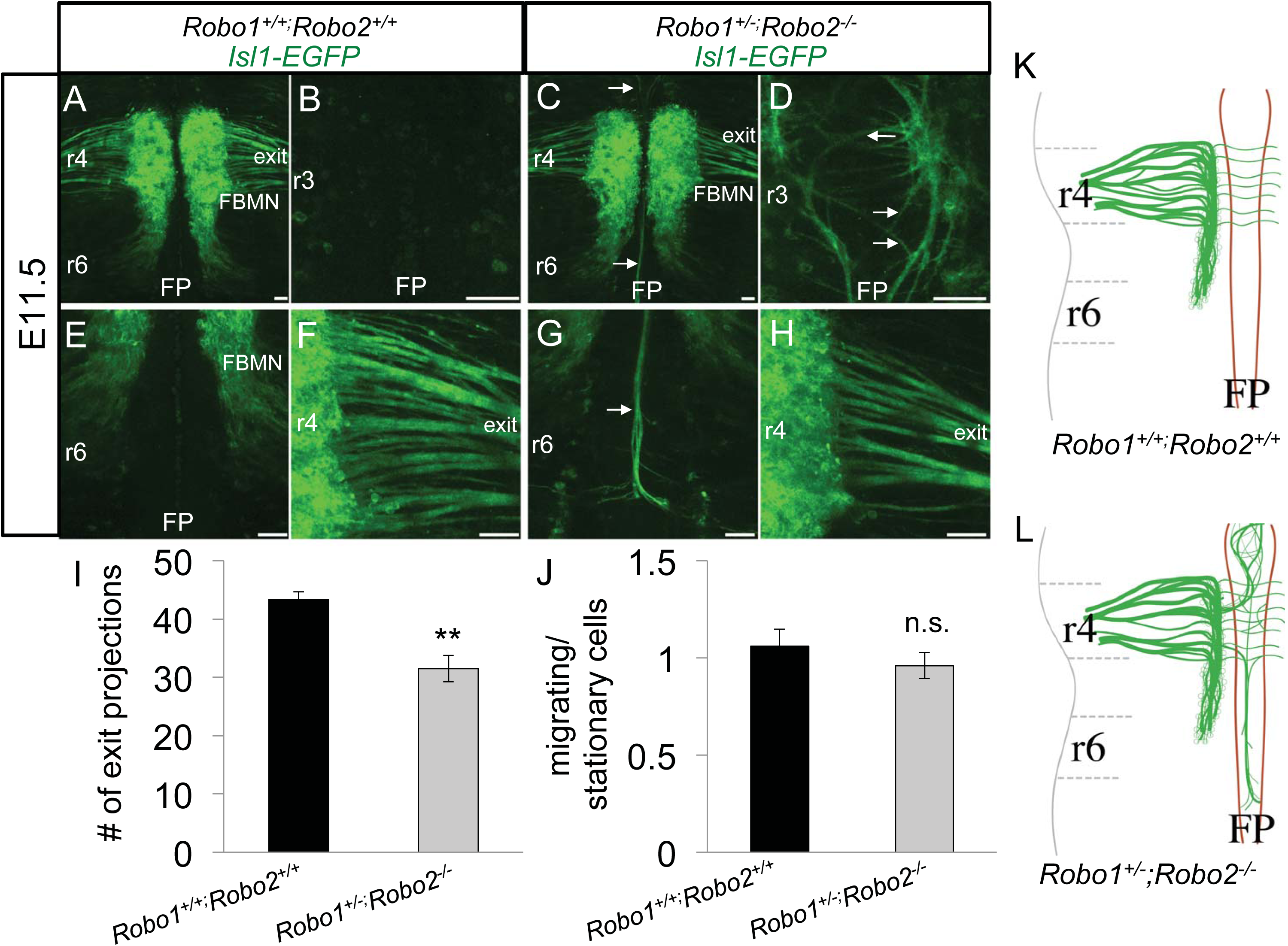
Robo1 is redundant with Robo2 in FBMN axon guidance. E11.5 open-book Isl1-EGFP hindbrains comparing control **(A-D)** and *Robo1*^*+/−*^;*Robo2*^*−/−*^ **(E-H)** embryos. **(A, D)** Control FBMN nuclei undertook caudal migration towards r6 and project axons to r4 exit point. **(E-G)** *Robo1*^*+/−*^;*Robo2*^*−/−*^ animals displayed more dramatic collapse of FBMN axons into the floor plate. **(F)** Axons from r4 were observed looping around in the FP of r3. **(G)** Multiple axons migrated caudally into the r6 floor plate in a large bundle. **(H)** Many but not all axons projected successfully towards the r4 exit point. **(I)** Measurement of the number of fascicles successfully navigating towards the r4 exit point in control and *Robo1*^*+/−*^;*Robo2*^*−/−*^ embryos. **(J)** Quantification of the ratio of migrating (r5-r6) cell bodies divided by stationary cell bodies (r4). **(K, L) (K)** Schematic of control E11.5 morphology compared to **(L)** *Robo1*^*+/−*^;*Robo2*^*−/−*^ mutants, which had increased numbers of axons migrating into the FP. Scale 50 μm. n=3, E11.5 *Robo1*^+/+^;*Robo2*^+/+^. n=4 (I)/n=3(J), E11.5 *Robo1^+/+^;Robo2^−/−.^*

### Robo1 and Robo2 guide initial facial axon outgrowth towards the exit point

To fully test if Robo1 and Robo2 act together to guide the FBMN, we examined the FBMN development of *Robo1*^*−/−*^; *Robo2*^*−/−*^ double mutant embryos, and also looked throughout the time course of FBMN development from E10.0-13.5. On E10.0 in control Isl^MN^;GFP embryos, facial cell bodies differentiated adjacent to the floor plate in r4 (Figure 3A). These cells had already sent pioneer axons to their ipsilateral dorsal motor exit point in r4 (Figure 3B). Some of the initial projections to the exit point had already formed thick bundles, however there were still individual axons not yet fasciculated. On E10.0, facial cell bodies were stationary along the ventral aspect of r4, and had not yet begun migrating caudally.

**Figure 3.**
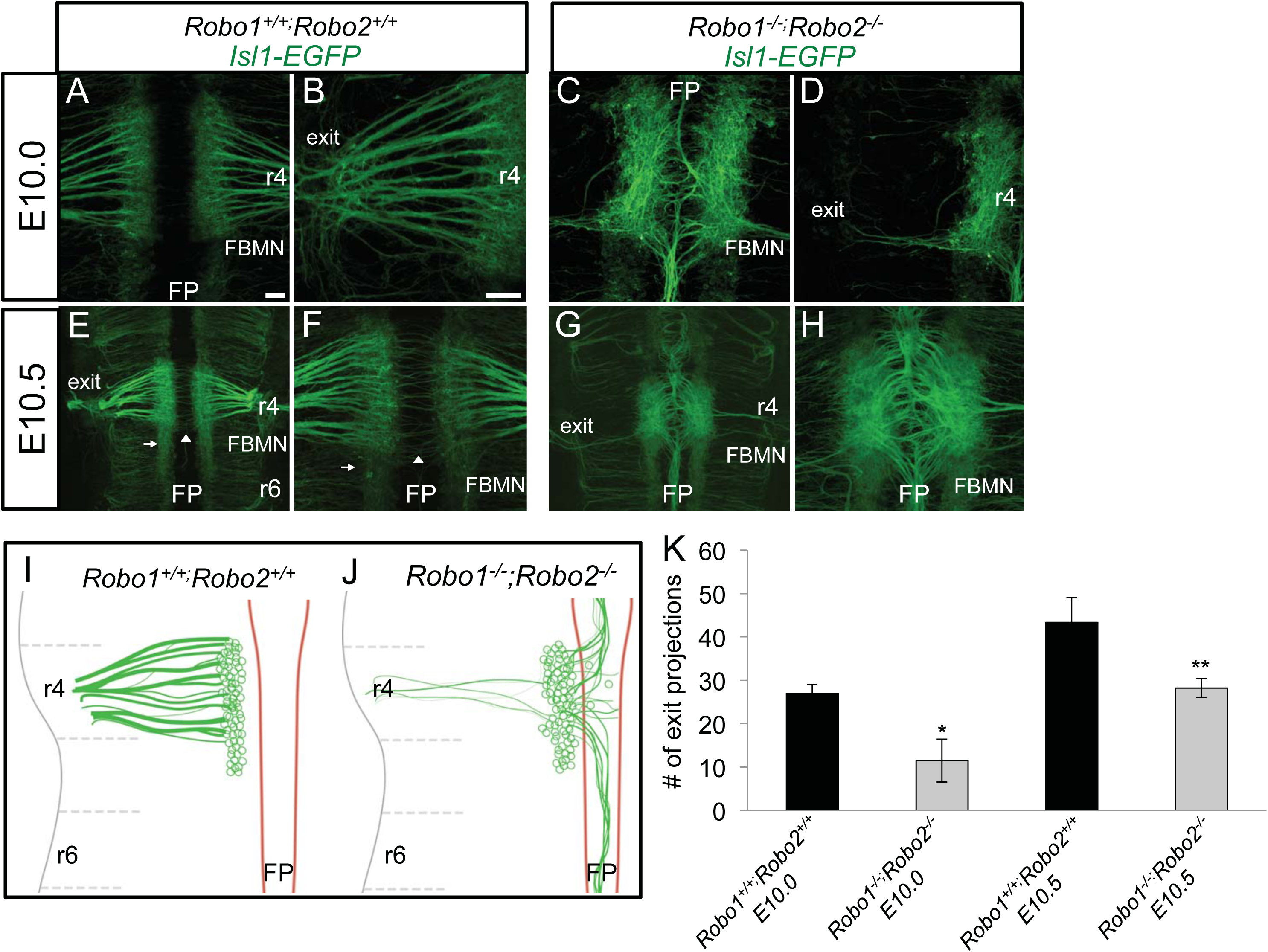
Robo1 and Robo2 are required to guide pioneer facial axons toward their r4 dorsal exit point. E10.0-E10.5 open-book Isl1-EGFP hindbrains comparing control (A, B, E, F) and *Robo1*^*−/−*^;*Robo2*^*−/−*^ (C, D, G, H) embryos. (A-D) E10.0 mouse hindbrains arranged rostral, top; floor plate (FP), center. (A, B) In control embryos, facial axons coursed from the facial branchiomotor nucleus (FBMN) towards the dorsal motor exit point (denoted as exit) in r4; a higher magnification view is shown in panel B. (C) In *Robo1*^*−/−*^;*Robo2*^*−/−*^ embryos, motor axons were misguided into the floor plate. Likewise, the nucleus was shifted closer to the floor plate. (D) A higher magnification view in *Robo1*^*−/−*^;*Robo2*^*−/−*^ mice to show that few axons projected dorsally from the FBMN to their exit point. (E-H) By E10.5, facial axons did not self correct after initial projection errors. (E,F) On E10.5 in control, FBMN cell bodies migrated caudally (arrow). The commissural axons of the inner ear efferents (IEE) nerve (arrowhead) also crossed the midline. (G) E10.5, facial axons did not project towards the exit point and instead projected in bundles rostrally and caudally in the floor plate. (H) The nucleus was wider and closer to the floor plate relative to control. (I, J) Schematics of E10.0-10.5 FBMN nerve projections. In control many axons projected to the exit point in rhombomere 4 (I). In Robo1/2 mutants, axons collapsed into the floor plate and nuclei shifted towards the floor plate (J). The tangential caudal migration of cell bodies started on E10.5, and was characterized by a subpopulation of FBMN pioneering this route. (K) Graph quantifying the number of facial axon bundles projecting to the r4 exit point in E10.0 and E10.5 control (black) compared to *Robo1*^*−/−*^;*Robo2*^*−/−*^ mutants (diagonal). Scale 50 μm. Error bars show S.E.M.; significance was measured using students t-test, *p<0.05; **p<0.01. n=3, E10.0 *Robo1*^+/+^;*Robo2*^+/+^. n=2, E10.0 *Robo1*^*−/−*^;*Robo2*^*−/−*^. n=3, E10.5 *Robo1*^+/+^;*Robo2*^+/+^ E10.5. n=5, *Robo1^−/−^;Robo2^−/−.^*

In *Robo1*^*−/−*^; *Robo2*^*−/−*^ Isl^MN^;GFP E10.0 embryos, severe facial axon and cell body guidance defects were already evident. The majority of pioneer facial axons projected into the floor plate in mutant animals (Figure 3C, K). Bundles of axons coursed through the floor plate both rostral and caudal relative to r4. On E10.0, these misguided axons did not stall, but migrated long distances in the floor plate. Correspondingly, the number of axons reaching the dorsal motor exit point in r4 was significantly reduced by 57% in *Robo1*^*−/−*^; *Robo2*^*−/−*^ animals (Figure 3D, K), which was more dramatic than that observed in *Robo1*^+/−^; *Robo2*^−/−^ animals (Figure 2I). Interestingly, the distance between the nuclei was also reduced in *Robo1*^*−/−*^; *Robo2*^*−/−*^ embryos, suggesting that facial neurons shifted toward the ventral midline, consistent with previous observations that cell body position is regulated by Robo signaling repelling motor neuron cell bodies from the midline (Kim et al., 2014).

On E10.5 in controls, facial cell bodies started migrating tangentially from r4 to r6. This rostral to caudal migration was initiated by a few pioneer cell bodies that reached r5 at this stage (Figure 3E). Additional differentiated facial cell bodies appeared as the nucleus grew larger in r4. Axons that coursed to the exit point in r4 appeared more tightly fasciculated at this stage. On E10.5, the initial outgrowth of the IEE commissural axons crossed the floor plate at r4 (Figure 3F).

On E10.5 in *Robo1*^*−/−*^;*Robo2*^*−/−*^ double mutant embryos, FBMN defects were much more severe than in single mutants. Some facial cell bodies initiated the expected tangential caudal migration in mutant animals (Figure 3G), though IslMN:GFP was expressed in all developing somatic motor neuron populations at this stage making the facial nucleus difficult to discern among the numerous anterior motor neurons present. Axons did not appear to correct their guidance errors from E10.0. The width of fascicles in the floor plate increased, perhaps due to additional axons that use pioneer axons already in the floor plate as a scaffold to migrate onto (Figure 3H). Some new axons also migrated onto bundles that already correctly exited the hindbrain into the periphery, but did not appear to forge a new pathway to the motor exit point. Interestingly, short neurites that projected off the cell bodies in r4 were present on this stage, which were not observed in control controls, suggesting growth cone stalling also occurs in the absence of Robo1/2.

### Robo1^−/−^;Robo2^−/−^ facial cell bodies make several types of migration errors during tangential migration

To observe the tangential migration in control and mutant animals we again utilized the Isl^MN^;GFP transgene on E11.5, and then used lipophilic dye tracing to follow their migration trajectories in E12.5 embryos. In controls, facial cell bodies started migrating caudally towards r6 on E11.5 (Figure 4A). Some cell bodies were still present in r4 on this stage, however the majority of cell bodies were migrating caudally (Figure 4B, O). Some leading cell bodies arrived in ventral r6 by this stage, though in this early stage of rostral-caudal migration, many cell bodies were still translocating through r5. In the process of moving, cell bodies laid down trailing axon tracts indicative of their migratory route. This created a mixture of cell bodies and motor axon tracts in ventral r4-r6 at this stage. In *Robo1*^*+/+*^; *Robo2*^*+/+*^ control animals, the separate IEE commissural axons had already traversed across the floor plate on this stage (Figure 4C). There were a few longitudinal axons visible in the embryonic floor plate, indicating a low level of errors attributed to the stochastic nature of axon guidance.

**Figure 4.**
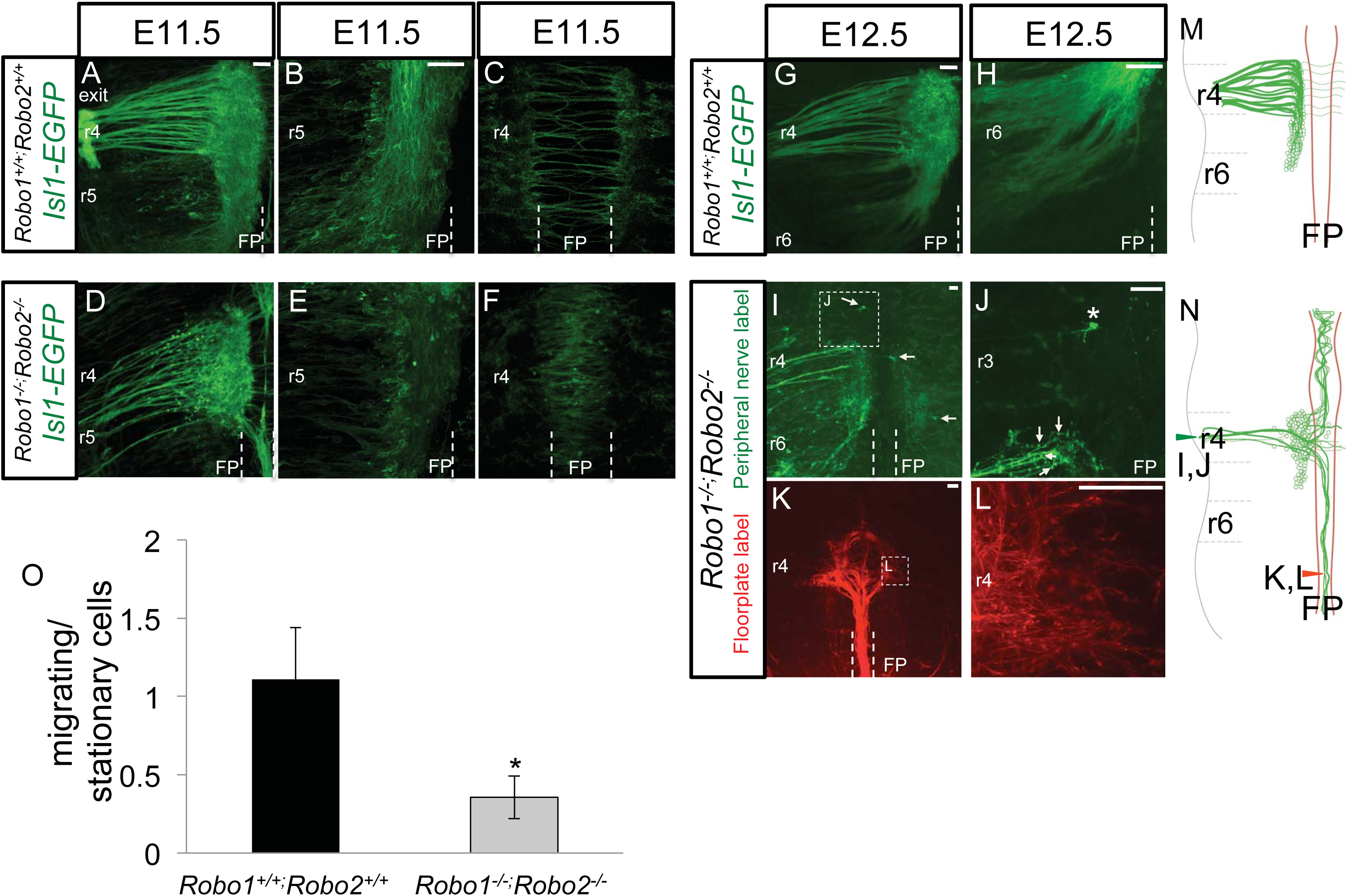
Robo1 and Robo2 regulate facial branchiomotor cell body position. Control **(A-C, G, H)** and *Robo1*^*−/−*^;*Robo2*^*−/−*^ **(D-F, H-L)** E11.5 and E12.5 open-book hindbrains expressing Isl^MN^:GFP or labeled with NeuroVue (Orange), respectively. **(A-I)** Mouse hindbrains arranged as in Fig 1. **(A)** On E11.5, facial axons added to fascicles already projecting to the motor exit point in control embryos. **(B)** FBMN cell bodies migrate tangentially from ventral r4 towards r6. The majority of cell bodies were in r5, however some cell bodies had already made it to r6 by E11.5. **(C)** Commissural axons of the IEE were the only axons seen crossing the floor plate in control embryos. **(D)** In mutant E11.5 animals, FBMN cell bodies were ectopically positioned more dorsally in r4. In *Robo1*^*−/−*^;*Robo2*^*−/−*^ embryos, axons added to the existing pioneer axons projecting to the exit point or floor plate. The fascicles coursing through the floor plate had increased in size relative to E10.5 embryos, suggesting that some newly projecting axons did not find the r4 exit point and instead followed axons into the floor plate. **(E)** There were significantly less cell bodies migrating caudally in mutant embryos, with the majority still residing in r4 (quantified in O). **(F)** On E11.5, cell bodies were ectopically located in the floor plate. **(G)** By E12.5, facial axons projecting to the exit point were tightly fasciculated at r4, and had laid down axon tracts tracing the cell bodies migration to r6. **(H)** Cell bodies largely completed their journey to r6. (**I, J)** To visualize facial axons, or aberrant axons that also exited r4 motor exit point, lipophilic dye was applied in the periphery (as described in methods). On E12.5 mutant embryos facial cell bodies appeared mispositioned (arrow) and disorganized. **(I)** Axon tracks spanning ventral r4 to r6 laid down from the migrating cell bodies appeared disorganized. Additionally, cell bodies from the contralateral side projected axons to the opposite exit point. **(J)** A magnified view of **I** demonstrating a cell body (asterisk) in r3 that had either sent an axon to the wrong exit point in r4, or had itself become misguided rostrally in r3. Ectopic facial cell bodies (arrows) in r4 were visible using lipophilic back labeling **(K, L)** *Robo1*^*−/−*^;*Robo2*^*−/−*^ E12.5 hindbrains were labeled with lipophilic dye in the floor plate of r8 to trace fascicles coursing caudally in the floor plate. **(K)** Back-labeling neurite bundles in the floor plate revealed these originate in the facial nucleus at r4. **(L)** *Robo1*^*−/−*^;*Robo2*^*−/−*^ cell bodies maintained long neurites in the floor plate. These same cell bodies also produced axon-like filaments heading towards the motor exit point, or stalling closely to the soma. **(M, N)** Schematics of the rostral-caudal migration of the FBMN cell bodies, as well as the nerve morphology, that occurs from E11.5 to E12.5 in both control **(M)** and *Robo1*^*−/−*^;*Robo2*^*−/−*^ **(N). (M)** In control embryos the majority of cell bodies have migrated to r5 and r6. Concurrently, newly differentiated cell bodies sent out axonal projections that adhere and migrate unto the pioneer tracts that had went through the motor exit point. The IEE axons were still the only axons in the floor plate. **(N)** In *Robo1*^*−/−*^;*Robo2*^*−/−*^ mutant facial cell bodies stalled in r4 and did not have as many migrating cells as controls. A subpopulation of cell bodies that remained in r4 were mispositioned dorsally, some of which possess multiple neurites. Newly projecting axons thickened the already misguided fascicles that continued to migrate rostrally and caudally within the floor plate. **(O)** Bar graph quantifying the proportion of migrating cell bodies (present in r5 and r6) over the number of stationary cell bodies (present in r4) in E11.5 embryos (area of cell bodies in r5 and r6 and divided by the area of cell bodies in r4). It was found using a student’s t-test that there are significantly (p=0.0104) more cell bodies migrating caudally in control (black) compared to mutant (diagonal) FBMN. Scale bar 50 μm. Error bars show S.E.M. *p<0.05 n=5 control, n=3 *Robo1^−/−;^Robo2^−/−.^*

In *Robo1*^*−/−*^; *Robo2*^*−/−*^ E11.5 facial nuclei, migratory defects of the cell bodies were apparent. Ectopic cell bodies were seen along the axon bundles projecting towards the exit point, thus showing a highly abnormal dorsal migration in r4 (Figure 4D). The nucleus also appeared larger in r4 in comparison to controls, suggesting that cell bodies stalled in r4 rather than migrating caudally as normal. In agreement with cell bodies not initiating migration in r4, there were significantly less cell bodies in mutant embryos that had migrated caudally to r5 compared to controls (Figure 4E, O). This is in contrast to *Robo1*^+/−^; *Robo2*^−/−^ animals, which did not have cell body migration defects (Figure 2J). Despite there being fewer cell bodies in r5, those that started migrating along the rostral– caudal axis appeared to successfully do so and migrated to r6 with the same timing as in controls. Surprisingly, some cell bodies also had migrated into the floor plate, presumably following their mis-guided axons to join the thickened longitudinal fascicles already there (Figure 4F).

On E12.5, to better determine the morphology of the FBMN that successfully migrate to the periphery, as well as to validate the identity of the longitudinal axons in the floor plate, dye-tracing experiments were performed in both control and mutant embryos, though only mutant labels are shown. Based on IslMN:GFP visualization of the FBMN (Figure 4G,H), and retrograde labeling from the peripheral nerve (data not shown), on E12.5 the rostral-caudal migration is mostly completed in control embryos. Exit point fascicles merged into large bundles on this stage (Figure 4G). In *Robo1*^*+/+*^; *Robo2*^*+/+*^ control embryos, most cell bodies were seen in r6 (Figure 4H). Though tangential migration was nearly complete on E12.5, we noted some cell bodies were still in r5. The presence of cell bodies en route to r6 was confirmed in both Islet-GFP embryos and dye tracing experiments.

To visualize the morphology and number of cell bodies that had successfully projected axons out to their exit point, retrograde tracing with lipophilic dyes were placed on either side of the peripheral facial nerve (Figure 4I, J). The migratory path of the cell bodies toward r6 appeared disorganized as evidenced by the axon tracts that followed the cell bodies spanning r5-r6 (Figure 4I). Ectopic cell bodies located rostral to r4 or within the floor plate of r4 were seen in E12.5, similar to E11.5 (Figure 4I, J). Retrograde dye labeling experiments revealed axon guidance errors that were not as evident in IslMN;GFP embryos. Retrograde labels from the exit points labeled mostly ipsilateral cells, as expected, but FBMN cell bodies contralateral to the dye label site were also labeled (Figure 4I). Contralateral neurons could be labeled if either contralateral neurons initially projected across the floor plate to the incorrect motor exit point, or if cell bodies migrated contralaterally across the floor plate. Other abnormal axon behaviors such as looping around in the floor plate were also clearly seen. Interestingly, a small set of cell bodies produced multiple long neurites (Figure 4I, J). Multipolar cell bodies labeled in this experiment possess an axon that successfully made it out to the exit point, as evidenced by their being labeled by dye placed in the periphery, and also another neurite that extended long distances into the floor plate. While control facial cell bodies normally extend small axon-like leading processes to guide tangential migration, these never extended as long as in *Robo1*^*−/−*^; *Robo2*^*−/−*^ animals.

To confirm that longitudinal tracts in the floor plate had indeed came from the facial nucleus, floor plate dye labels more rostral (r1) and caudal (r8) relative to r4 were performed (Figure 4K, L). Back-labeled axons were traced back to cell bodies stalled in r4 (Figure 4K), or to cell bodies in r6 (data not shown). The majority of cell bodies in r4 possessed short growth cones, which implies that, in addition to their primary but mis-guided long axons, they were producing additional neurite outgrowths. Despite possessing long neurites that coursed longitudinally through the floor plate, some cell bodies back-labeled to r4 also possessed filaments that extended towards the r4 motor exit point, consistent with the retrograde labels from the exit points (Figure 4L).

### Facial Branchiomotor cell bodies extend multiple neurites in Robo1/2 double mutants

By E13.5 in control embryos, facial cell bodies and corresponding axons completed their migration through the hindbrain. To specifically back label and visualize the facial nucleus, and to avoid the over-crowding seen in Isl1-GFP labels, lipophilic dye was placed on the peripheral facial nerve (Figure 5A-H). Cell bodies had completed their migration along the tracts of facial axons laid down between r4-r6 (Figure 5A). Axon bundles exiting in r4 had tightly fasciculated at this stage. Immediately after E12.5, the facial nuclei started migrating away from the ventral location to a new dorsal-lateral position, still in r6. Concurrently, cell bodies also migrated radially away from the ventricular surface to the pial surface of r6 (Figure 5B). In control embryos, the only axons in the floor plate or contralateral side were those of the IEE population (Figure 5E, F).

**Figure 5.**
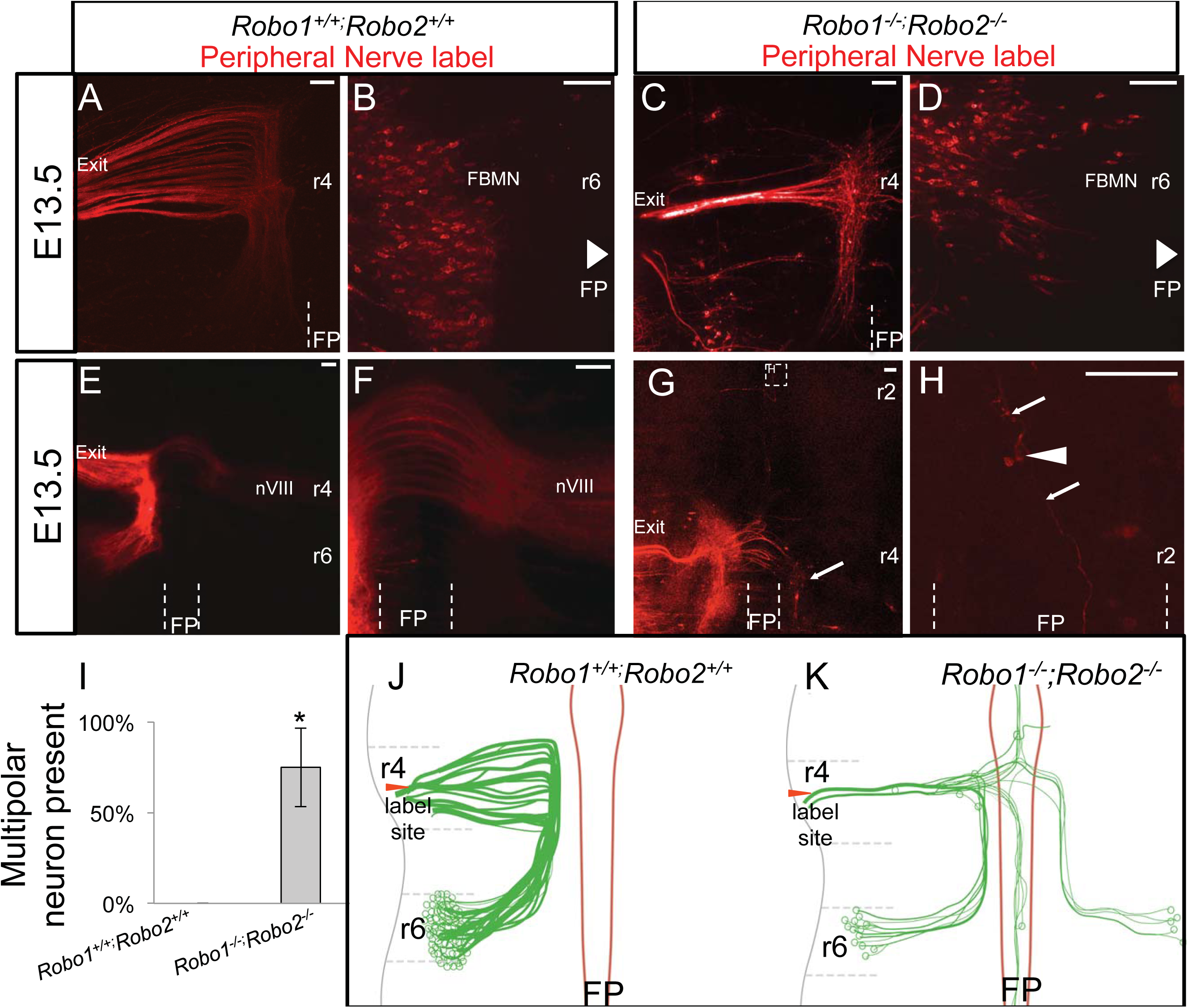
Subpopulations of facial cell bodies extend multiple long neurites in the absence of Robo1/2. E13.5 open-book *Robo*^*+/+*^;*Robo2*^*+/+*^ (A,B,E,F) and *Robo1*^*−/−*^;*Robo2*^*−/−*^ **(C, D, G, H)** facial nuclei back labeled with NeuroVue Orange dye. **(A-H)** E13.5 mouse hindbrains arranged rostral, up; floor plate (FP), center. Embryos imaged ventricular side up **(A, C, E-H)** or pial side up **(B,D)**. **(A-D)** Demonstration of the final dorso-lateral and radial migration patterns in both control and mutant embryos. **(A, B)** *Robo*^*+/+*^;*Robo2*^*+/+*^ facial axons exited in r4 **(A)**, while the cell bodies **(B)** radially migrated to the dorsolateral pial surface of r6. **(C)** Axons in *Robo1*^*−/−*^;*Robo2*^*−/−*^ animals were reduced in number and appeared disorganized in the rostral caudal tract. **(D)** Cell bodies that correctly migrated rostral-caudally in Robo1/2 double mutants also successfully migrated to the pial surface of r6. **(E-H)** Illustrates the axon migration patterns of control and mutant embryos in a late stage. **(E, F)** *Robo*^*+/+*^;*Robo2*^*+/+*^ facial nerves followed their axons to the lateral side of r6. The contralateral projections of the IEE were the only axons seen in or across the floor plate. **(G)** E13.5 *Robo1*^*−/−*^;*Robo2*^*−/−*^ facial axons migrated long distances incorrectly throughout the hindbrain. Cell bodies of the contralateral side incorrectly sent out axons (arrow) to the exit point of the opposite side. Cell bodies extended axons to exit point and also a separate neurite into the floor plate. Cell bodies with neurites in the floor plate as well with projections in the floor plate were observed in 3/4 *Robo1*^*−/−*^;*Robo2*^*−/−*^ embryos and 0/6 in *Robo^+/+^;Robo2^+/+^*. **(H)** A magnified view of a cell body (arrowhead) in the floor plate that produced two long neurites (arrows). **(I)** Percent embryos with observed multipolar neuron in control and Robo1/2 mutant embryos. **(J, K)** Schematics of E13.5 FBMN nerve projections demonstrating the dorsolateral and radial migrations of the cell bodies and axons that followed. **(J)** In control, axons followed the migration path of the cell bodies to the pial surface of r6 while bundles continued to fasciculate on their way to the r4 exit point. Cell bodies were arranged lengthwise across the entire pial surface of r6. **(K)** In *Robo1*^*−/−*^;*Robo2*^*−/−*^ animals, cell bodies sent axons out to the contralateral exit point and were sometimes ectopically located in the floor plate. A subpopulation of cell bodies also extended long and multiple neurites.

In *Robo1*^*−/−*^; *Robo2*^*−/−*^ double mutant embryos, although the nerve was much smaller, some axons could be back-labeled from the nerve exit point. The retrograde label successfully traced peripheral axons back to their cell bodies, but also labeled other long neurites that migrated abnormally. A small subpopulation of the back-labeled FBMN migrated successfully to the correct location. However, many mutant cell bodies were displaced. Tracks of axons spanning r4 to r6 were disorganized, much like what occurred on E12.5 (Figure 5C). Axon tracts that successfully arrived at the exit point appeared to merge into one or two bundles that exited the hindbrain. Radial migration appeared largely unaffected in those mutant neurons that did reach r6, as some cell bodies reached the pial surface of r6 (Figure 5D). Similarly, cell bodies correctly located on the pial surface were also in the correct dorsal-lateral position (Figure 5D). Stalling cell bodies were still present in r4 and r5 in mutant embryos (Figure 5C), and we noted that these cell bodies were not at the pial surface (not shown). Additionally, ectopic cell bodies migrated along the axon bundles reaching the r4 motor exit point. Axon guidance defects within the hindbrain remain prevalent at this stage as well. Because these E13.5 labels were retrograde from the exit point, many of the long axons projecting into ectopic tracts apparently represent second (or multiple) axons from FBMN neuron cell bodies. *Robo1*^*−/−*^; *Robo2*^*−/−*^ embryos had extensive neurons and processes growing into the floor plate, including longitudinal tracts, looping axons, and cell bodies (Figure 5G, H). Facial cell bodies on the contralateral side of the dye site extended axons to the wrong contralateral exit point (Figure 5G). Cell bodies located in other rhombomeres also incorrectly sent axons to the r4 exit point and possessed multiple long neurites (Figure 5H). Therefore, the E13.5 labels show an accumulation of migration errors of both cell bodies and axons, and reveal additional examples of FBMN neurons producing two or more long axonal projections.

## Discussion

In our study, we found that Robo1/2 has a multifunctional role in regulating both the facial nucleus migration and the motor exit point pathfinding. We established that Robo1 and Robo2 receptors function redundantly to guide both FBMN axons and cell bodies. Examining *Robo1*^*−/−*^; *Robo2*^*−/−*^ embryos revealed much stronger and more diverse phenotypes than observed in Robo1 and Robo2 single mutant embryos or even *Slit1*^*−/−*^; *Slit2*^*−/−*^ double mutant animals (Hammond et al., 2005). First, in *Robo1*^*−/−*^; *Robo2*^*−/−*^ many facial motor axons fail to make it to the r4 motor exit point and instead divert into the floor plate, which implies Slit proteins secreted from the floor plate (Holmes et al., 1998; Yuan et al., 1999) act to repel facial axons dorsally towards their exit point. Additionally, as facial axons collapse into the floor plate they are likely attracted to floor plate secreted cues such as Wnt5a and Wnt7a (Vivancos et al., 2009). Secondly, the caudal migration of the nucleus is disrupted in Robo1/2 mutants, as cell bodies are observed stalling in r4 or migrating incorrectly into regions such as the floor plate, thus suggesting Robo-Slit signaling is required to guide facial cell bodies during tangential migration to r6. Lastly, we observed multiple long neurites projecting off facial cell bodies during its migration, suggesting Robo is important for suppressing the formation of ectopic or secondary neurites from FBMN neurons.

Numerous motor neuron populations express Robo1 and Robo2 and are responsive to Slit2 (Bravo-Ambrosio and Kaprielian, 2011; Brose et al., 1999; Holmes et al., 1998; Jaworski et al., 2010), including the branchiomotor aspect of the facial nerve and nucleus (Hammond et al., 2005; Murray et al., 2010). All three Slits (Slit1, Slit2, Slit3) are highly expressed in the floor plate, spanning the length of the midbrain through the spinal cord (Brose et al., 1999; Holmes et al., 1998; Yuan et al., 1999). Slit2 is expressed in the ventral hindbrain during the entirety of migration of the facial nucleus, from E8.5 before facial cell bodies are even differentiated through E14.5 when the nucleus has settled to its final location (Geisen et al., 2008). Motor neurons also express Slit2 and 3, potentially regulating themselves in an autocrine fashion (Jaworski and Tessier-Lavigne, 2012). Interestingly, at the facial nucleus itself expresses Slits (Geisen et al., 2008; Kim et al., 2016), suggesting the facial nucleus may regulate its guidance or organization at later stages of development, a potential guidance mechanism that could be tested by cell-type specific knockouts.

### Facial nerve projections towards the motor exit point are guided by Robo-Slit signals

Our study validates that Robo-Slit signaling guides branchiomotor axons out of the floor plate. In our study, *Robo1*^*−/−*^; *Robo2*^*−/−*^ double mutants exhibited more severe defects than single Robo1 or Robo2 or *Slit1*^*−/−*^; *Slit2*^*−/−*^ double knockouts (Hammond et al., 2005). Our extreme phenotypes with the axons, and cell bodies alike, suggests Robo1 and Robo2 largely act redundantly to regulate this population. The previous group had found that some mutant axons migrated the floor plate and also crossed between the exit point fascicles in Robo mutants (Hammond et al., 2005). Agreeing with these results, our Robo1/2 double knockout animals we found most axons of the facial nucleus migrate into the floor plate. The *in ovo* electroporation of a dominant-negative Robo receptor experiments performed by Hammond et al. strongly agree with the phenotypes we saw in our *Robo1*^*−/−*^; *Robo2*^*−/−*^ animals. They found axons that stalled, looped around, and failed to project away from the floor plate and to the exit point in their chicken experiments, which are all effects we saw in our mutant animals. The facial axons collapsing into the floor plate are largely seen migrating rostral or caudal relative to r4, suggesting there are attractive cues guiding them to these locations. In agreement with this, it was previously found Wnt5a and Wnt7a act as chemoattractants for facial axons (Vivancos et al., 2009). Indeed, Wnt5a and Wnt7a are expressed in the floor plate rostral and caudal to r4, respectively, and thus may be attracting facial axons that are no longer repelled by Robo-Slit signaling (Vivancos et al., 2009). In another study, the chemorepulsive effects of the Robo ligand Slit, and the Unc5a ligand Netrin, were previously both found to repel facial axons (Murray et al., 2010). Netrin was found to be particularly important in guiding facial neurons to their dorsal exit point, as facial axons in Netrin mutants turn towards the exit point too early or migrate into the floor plate (Murray et al., 2010). This is very similar, though less severe, than what we see in Robo1/2 double mutants suggesting there may be redundancies or interactions between Netrin and Robo signals.

Slit-Robo signaling is important for motor exit point pathfinding. For instance Robo2, under the regulation of Nkx2.9, is necessary for spinal accessory motor neurons to correct exit the neural tube (Bravo-Ambrosio et al., 2012).The neural crest-derived motor exit point is joined later in development by boundary cap cells that keep cell bodies from leaving the periphery where at the same time are thought facilitate axon exit out of the neural tube by secreting an attractant (Niederländer and Lumsden, 1996). Boundary cap cells of the spinal cord produce various chemorepellents (Sema3B/3G/6A) that are thought to repel the chemoreceptor (Neuropilin-2, PlexinA1) expressing motor cell bodies away from the exit point (Bron et al., 2007; Mauti et al., 2007). Motor axons are not thought to express these chemorepulsive receptors, and are therefore permitted to migrate towards the motor exit point to leave the neural tube. In agreement is the finding, facial motor axons are not repelled by the motor exit point expressing Sema3A, but are instead repelled by Slit which appears to flank the exit point (Murray et al., 2010; Yuan et al., 1999). The expression pattern of Slit1/2 being adjacent to the motor exit points suggests Slits may be corralling axons to the correct exit point in the right rhombomere level. Together these data suggest Robo-Slit signaling is necessary for accurate guidance of the developing FBMN axons towards their dorsal r4 motor exit point.

### Caudal migration requires multiple signaling cascades, including Robo1/2

In our study, we found that Robo-Slit signaling is important for caudal migration of the facial nucleus. While it is well established that Slit-Robo signals can regulate dorsal-ventral migrations of axons, it is not obvious how the ventral Slits could affect anterior-posterior migration of cell bodies. One possibility is that Robo1/2 signaling integrates with other previously known guidance pathways to guide this complex migration of the cell bodies. This is in contrast to Hammond et al., who found Robo mutants only have axon defects (Hammond et al., 2005). This difference is most likely attributable to using Robo1/2 double knockout mice and previous work being done in single Robo1 or Robo2 knockouts. Our results are in agreement with experiments in which Hammond et al. found mispositioned cell bodies when electroporating dominant-negative Robo constructs into chick embryos (Hammond et al., 2005). We found that in Robo1/2 mutant mice, some facial cell bodies remain in ventral r4 and do not undergo tangential migration. A few mutant cell bodies migrate into the periphery along their axons, whereas others move into the floor plate. These abnormal migrations could indicate that Robo1/2 signaling either acts to instigate, allow, or modulate caudal migration. The PCP pathway is necessary for the initial migration of the facial nucleus. The PCP pathway guiding the facial nucleus functions in an atypical Disheveled (Dvl) independent manner (Glasco et al., 2012), which is normally an essential component of most Wnt pathways (Gao and Chen, 2010). Our results indicate Robo-Slit may integrate with the PCP pathway to mediate the caudal migration of the facial nucleus; further research is necessary to examine this potential integration of Robo-Slit signaling and the PCP pathway.

The migration defect in Robo1/2 mutants does not occur in all mutant facial cells nor are cell bodies mispositioned in a consistent way, making it difficult to ascertain how Robo1/2 is regulating cell body position. One mechanism by which Robo-Slit signaling could affect tangential migration is by orienting leading processes away from, or parallel to, the floor plate, which is apparently highly attractive in the absence of Robos. It is also possible that the Slit-expressing dorsal neural tube also has a repulsive influence, thus restricting migration in a longitudinal direction.

One possible attractive receptor and ligand complex that Robo-Slit signaling may modulate to orient cells caudally is Frizzled (Fz) and Wnt. Wnt5a and Wnt7a attract migrating facial neurons, however when these molecules are removed the nucleus is mostly unaffected (Vivancos et al., 2009). When inhibiting the Wnt5a canonical effector proteins ROCK, MLCK, and JNK, the migration of the FBMN was more severe than in Wnt5a knockouts alone (Vivancos et al., 2009). This suggests that other axon guidance receptors, such as Robo, also redundantly activate these effectors to guide the facial nucleus. In agreement with this idea, it has been shown Slit activates similar downstream effectors as Wnt5a-MLCK, ROCK, and Myosin II—to repel facial axons (Murray et al., 2010). Furthermore, as Dvl is not necessary for migration of the nucleus (Glasco et al., 2012), but is normally required downstream of Wnt signaling to activate GTPases that activate MLCK, ROCK, and MyosinII (Rosso et al., 2005; Strutt et al., 1997), perhaps Robo1/2-Slit are necessary to activate these GTPases in order for Fz-Wnt signaling to polarize the neuron outgrowth. This possibility is feasible due to Slit1/2 being expressed during the entirety of the facial nucleus migration in the floor plate adjacent to the tangentially growing cell bodies (Hammond et al., 2005; Holmes et al., 1998). It would be interesting to test the effect of knocking out both Robo1/2 and Wnt5a to see if these redundantly guide the facial nucleus. Another possibility is Robo signaling may also promote an asymmetric localization of other receptors necessary for the correct growth orientation, as suggested in nematode (Tang and Wadsworth, 2014).

Another potential mechanism Robo1/2 may be necessary for tangential migration is that it may attenuate cell adhesion at the start of migration, thus acting as a permissive signal. In *Robo1*^*−/−*^; *Robo2*^*−/−*^ mutants, we observed facial cell bodies that appear to be stalled in r4 through the time period of E11.5 to E13.5, thus suggesting the adhesion between the neuroepithelium and the facial nuclei are not alleviated to release the cell bodies to migrate. Robo regulates cell migration by attenuating cadherin adhesion, thus breaking apart junctions between cells (Rhee et al., 2007). During migration of the FBMN Robo may act to inhibit N-Cadherin mediated adhesion, as demonstrated in other systems (Rhee et al., 2007). Indeed, in zebrafish N-Cadherin is necessary for preventing facial cell bodies from entering the floor plate and is expressed in the surrounding neuroepithelium (Stockinger et al., 2011). The potential role of Robo1/2 signaling in mediating cell adhesion of the FBMN is an interesting possibility that warrants future study.

Lastly, the facial nucleus migration errors may be secondary or indirect effects of other substrates that were misplaced in Robo1/2 mutants. In our study, we observed cell bodies misguided into the floor plate, which may be due to them following other misguided populations to the floor plate. In zebrafish, facial neurons undergoing tangential migration through r5 and r6 use the medial longitudinal fasciculus (MLF), an ipsilateral tract of axons connecting visual and auditory nuclei of the brain to the spinal cord, as a substrate to migrate on (Wanner and Prince, 2013). Without the MLF, non-pioneer facial cell bodies did not migrate successfully to r6. Though it has been suggested that the FBMN is located too dorsally in mouse to contact the MLF (Wanner et al., 2013), this has not been experimentally validated. Our previous work has shown without Robo-Slit signaling, the MLF collapses into the floor plate and forms long fascicules coursing through the floor plate (Farmer et al., 2008; Kim et al., 2014). It would agree with our results if facial cell bodies adhered to the MLF as a substrate to mislead them into the floor plate. Alternatively, it is possible that facial axons may make independent ventral errors, because of the lack of Slit-mediated floor plate repulsion, but turn longitudinally once they contact the ectopic MLF bundles within the midline.

### Dorsolateral and radial migration act independently of Robo signaling

When facial cell bodies reach r6 on E12.5 they turn perpendicular towards the dorsal aspect of r6, a process complete on E14.5. Without Robo1/2 signaling, facial neuron cell bodies become misplaced during the period when caudal migration should be occurring and remain displaced in later stages. Though mis-positioned cell bodies do not appear to undergo dorsolateral migration on E12.5, it is not easy to determine if Robo is particularly important in regulating tangential migration as they are already strongly affected earlier in development. To ascertain whether Robo is needed in these later stages, an inducible or temporal specific knockout would be needed to see if the lack of tangential migration is due to earlier defects or a separate guidance mechanism.

In Robo1/2 double mutant mice, we observed the mutant cell bodies affected earlier in development do not migrate to the pial surface, though unaffected facial cell bodies do correctly migrating to the pial surface of r6. Similarly to the disruption of dorsolateral migration in Robo1/2 mutant cell bodies, it is difficult to know if the lack of ventricular to pial migration in cell bodies are due to earlier defects or are indeed important for radial migration. Though Robo doesn’t appear necessary for dorsolateral and radial migrations, the initial correct caudal location of the cell bodies may influence the ability of mutant facial cells to undergo dorsolateral migration.

### Robo1/2 signaling suppresses the formation of multiple long axons from facial motor neurons

In Robo mutant animals spanning E10.0-E13.5, we observed that some facial cell bodies produced multiple long projections. Through our retrograde labeling experiments of the peripheral nerve, we observed facial axons, labeled because they made long primary axons that successfully projected to the motor exit point, were connected to axons at the same time making long projections caudally in the floor plate. Previously, these ectopic longitudinal tracts were observed in relatively lower numbers in Robo2 mutants (Hammond et al., 2005), and were attributed to both the FBMN and the IEE, which normally send commissural axons across r4 (Fritzsch and Nichols, 1993). Though some of the longitudinal axons in the floor plate could be from the IEE in our mutant animals, our floor plate back labeling experiments on E12.5 showed that many longitudinal fascicles in the floor plate traced back to the location of the FBMN. Therefore, the retrograde labeling shows that Robo signaling has an unexpected function in suppressing the formation of two or more long axonal projections from motor neurons. In contrast, Slit/Robo signaling is known to promote sensory axon branching (Ma and Tessier-Lavigne, 2007; Ozdinler and Erzurumlu, 2002; Wang et al., 1999; Yeo et al., 2004), yet the loss of Robo in facial motor neurons appears to lead to long ectopic neurites. Moreover, it is surprising that motor neurons have the capacity to form multiple neurites, given their universal unipolar structure in vertebrates. The mechanism underlying Robo suppression of ectopic motor axons is unknown. However, it may be relevant to note that two of the rare examples of vertebrate motor neurons that migrate, the facial motor neurons and a subset of oculomotor neurons that cross the midline, both involve the formation of an axon-like leading process (Bjorke et al., 2016). We speculate that the ability of these subsets of motor neurons to form a secondary axon-like process must involve a repolarization of the cell body to generate a secondary growth center and projection, and evidently this repolarization is subject to Robo regulation.

Robo may regulate the length of neurites through activation of Cdk5. The kinase Cdk5 is well known to regulate neurite length (Ohshima et al., 1996, 2007; Tang and Wang, 1996). Some *Robo1*^*−/−*^; *Robo2*^*−/−*^ mutant facial nuclei successfully migrate to the pial surface of r6, indicating that Robo activation of Cdk5 may not be necessary for successful radial migration but for regulating neurite length and number. Robo-Slit signaling limits the number of neurites cell bodies extend (Quinn et al., 2006). Perhaps this may occur in part through Robo activating Abl and Cables (Bashaw et al., 2000; Rhee et al., 2007), the latter of which activates Cdk5 (Zukerberg et al., 2000). Cdk5 may in turn regulate the number of neurites projecting off cell bodies. A more simple mechanism may just be that Robo-Slit signaling normally repels these multipolar neurites out of the floor plate much like it repels longitudinal and post crossing commissural populations (Brose et al., 1999; Farmer et al., 2008; Long et al., 2004). Without this repulsive signaling coming from the floor plate, neurites and the cell body that follows may be attracted to the floor plate by unidentified cues, and continue extending as seen in other populations (Farmer et al., 2008).

## Experimental Procedures

### Mouse embryos

Animal protocols were approved by the University of Nevada, Reno’s Institutional Animal Care and Use Committee (IACUC) and in accord to the standards of the National Institutes of Health Guide for the Care and Use of Laboratory Animals. Embryonic mice were collected at embryonic day 10.0 (E10.0), E10.5, E11.5, E12.5, and E13.5. Noon of the day a plug was discovered was counted as E0.5. If between E10.0- E12.5 embryos were obtained through uterine dissection, put in fresh (made the same day) 4% paraformaldehyde (PFA) and fixed for a minimum of 24 hours. If embryos were E13.5 cardiac perfusion was performed. We bought CD-1 mice from Charles River Laboratory (Wilmington, MA). Limb buds were removed from freshly collected embryos to genotype Robo mutant embryos as previously described (Grieshammer et al., 2004; Plump et al., 2002). Robo1/2 founder mice were a generous gift of Marc Tessier-Lavigne, of Rockefeller University. The Isl1^MN^-EGFP strain was a gift of Samuel Pfaff (Lewcock et al., 2007), of the Salk Institute. To create stable heterozygous lines, Islet-GFP mice were crossed with Robo1/2 mutant mice as well as CD1 mice.

### NeuroVue facial nerve tracing

The lipophilic fluorescent axon tracers NeuroVue™Orange and NeuroVue™ Jade were used to label the facial nucleus and trace facial axon trajectories in the peripheral branchial arches (Fritzsch et al., 2005). To trace the facial nerve a small piece of dye was cut into a triangle shape, and inserted superior to the otic vesicle on E10.0 and E10.5.

Jade dye was placed above the left vesicle whereas orange was placed above the right vesicle. Similarly on E11.5, dye was anterior to the otocyst, and lastly on E12.5 and E13.5 dye was placed in the most rostral region within the developing ear. For peripheral nerve tracing experiments, *Robo*^*+/+*^; *Robo*^*+/+*^ and *Robo^−/−^; Robo^−/−^*, dye was placed in same location to assure consistency. For floor plate fascicle tracing experiments, a smaller piece of dye than used in peripheral tracing experiments was placed directly in the floor plate several rhombomeres caudally or rostral relative to r4. After dye was placed embryos were incubated for 24hrs (E10.0, E12.5), or 48 hrs (E13.5) at 37° C to allow the dye to diffuse to the distal nerve and nucleus.

### Imaging and preparations of tissue

Embryos were assessed for successful fluorescent labels under a Leica upright fluorescence microscope. Neural tubes were dissected out while preserving the peripheral branchial arch. Hindbrains were prepared in an “open-book” conformation by cutting the dorsal most aspect of the neural tube along the dorsal midline. Neural tubes were incubated in 80% Glycerol in 4% PFA for several hours before imaging. Once mounted, embryos were imaged using the Leica TCS-SP8 confocal microscope within 2 days of the end of dye diffusion to limited the amount of unwanted diffusion of the dye.

### Analysis and quantification

Images were analyzed using the NIH’s ImageJ software. Neural tube analysis: Number of exit point projections were analyzed by counting the number of dorsal projections coming immediately off the facial nucleus with all sizes of fascicles or axons were counted as equal. Length of fascicle in floor plate was measured by starting at the end of the IEE (r4/r5 boundary) and measuring the length of the fascicle caudally. Migrating cells were counted as the area facial cell bodies in r5 and r6, while stationary cell bodies were counted as the area of the facial nucleus cell bodies in r4. For E13.5 cell bodies were counted as multipolar if axons were labeled in the periphery and neurites also present in the floor plate. For figures 1-4 a two-tailed Student’s t-test was performed to determine significance; standard deviation was used to generate error bars. For figure 5 a N-1" Chi-squared test was performed to determine significance; standard error of the proportion was used to generate error bars.

## Acknowledgements

The *Robo1*^*−/−*^; *Robo2*^*−/−*^ mice were generous gifts from Marc Tessier-Lavigne (Stanford; Genentech). The Isl^MN^:GFP-F mice were kindly provided by Samuel Pfaff (Salk Institute). We would like to especially thank Bernd Fritzsch (University of Iowa) for his expertise and advice with dye labeling for this project, and for preliminary labels of cranial nerve exit points. We thank many members of the Mastick lab who provided feedback and help with the project, including Katherine Weller, Tatiana Fontelonga, Clare Lee, and Johnathan Pietz. This project was supported by NIH R01 NS054740, R21 NS077169, and R01 EY025205 to G.S.M. Research reported in this publication used the imaging core facility supported by the National Institute of General Medical Sciences of the National Institutes of Health under grant number P20 GM103440, P20 GM103554, and P20 GM103650.

